# A novel algorithm for the collective integration of single cell RNA-seq during embryogenesis

**DOI:** 10.1101/543314

**Authors:** Wuming Gong, Bhairab N. Singh, Pruthvi Shah, Satyabrata Das, Joshua Theisen, Sunny Chan, Michael Kyba, Mary G. Garry, Demetris Yannopoulos, Wei Pan, Daniel J. Garry

**Affiliations:** Medicine Department and the Lillehei Heart Institute, University of Minnesota, Minneapolis, MN 55455, USA; Division of Biostatistics, School of Public Health, University of Minnesota, Minneapolis, MN 55455, USA; Stem Cell Institute, University of Minnesota, Minneapolis, MN 55455, USA

## Abstract

Single cell RNA-seq (scRNA-seq) over specified time periods has been widely used to dissect the cell populations during mammalian embryogenesis. Integrating such scRNA-seq data from different developmental stages and from different laboratories is critical to comprehensively define and understand the molecular dynamics and systematically reconstruct the lineage trajectories. Here, we describe a novel algorithm to integrate heterogenous temporal scRNA-seq datasets and to preserve the global developmental trajectories. We applied this algorithm and approach to integrate 3,387 single cells from seven heterogenous temporal scRNA-seq datasets, and reconstructed the cell atlas of early mouse cardiovascular development from E6.5 to E9.5. Using this integrated atlas, we identified an Etv2 downstream target, *Ebf1*, as an important transcription factor for mouse endothelial development.

## Introduction

Single cell RNA-seq provides unprecedented opportunities to study the complex cellular dynamics during the entire embryonic developmental process by profiling the transcriptomes of single cells, from the zygote to the postnatal stage[1–9]. Due to the heterogeneity of underlying cell populations, separate studies usually focus on specific embryonic regions or subpopulations of cells that express specific gene markers, during a relatively narrow developmental window (e.g. several embryonic days). Such temporal scRNA-seq datasets are usually characterized by large variance between time points during the developmental process[10]. While each study discovers the states and cell types within a specific developmental context, the integration of the scRNA-seq data from independent studies will allow us to systematically reconstruct the temporal and spatial developmental trajectory, and comprehensively understand the entire developmental process.

One of the major technical challenges of joint analysis of multiple scRNA-seq datasets is that the scRNA-seq dataset generated at different times, by different laboratories, and/or using different experimental procedures, result in strong batch effects, where the gene expression difference between batches are as significant as that between distinct cell populations[11–13]. Thus, the biological variation in the joint scRNA-seq datasets from different studies is confounded by the data source. Although methods such as surrogate variable analysis[14] and RUV[15] have been widely used to correct the batch effects in analyzing the bulk RNA-seq data, Haghverdi et al. and Bulter et al. initially described computational approaches for the systematic correction of such batch effects in scRNA-seq datasets[12,13]. Several other algorithms have been proposed to improve the speed[16–18], to combine more diversified datasets without sharing a common subpopulation[16,19], to integrate multiple types of single cell data[20], or to fine-tune the resolution and breadth of the meta-analysis[21]. Principally, most of the batch correction methods look for batch-free low dimensional representation to enable shared cell type identification across datasets, by accounting for the systematic difference between batches.

Most of the existing batch correction methods adjust the batch effects by matching the similar cells or cell clusters between entire batches (e.g. by looking at the mutual nearest neighbors between all the cells from two batches)[12,16–19,21,22]. However, when applying this strategy to the temporal scRNA-seq data, the cell matching will be confounded by the time index, which usually has a strong correlation with the gene expression pattern[10]. This will result in the loss of the entire developmental trajectories on the joint low dimensional space.

Here, we developed a novel computational method for the integration of different sources of temporal scRNA-seq data, which we named single cell neighborhood component analysis (scNCA). We demonstrated that scNCA successfully integrated 3,387 single cells from seven heterogenous temporal scRNA-seq datasets of mouse early cardiovascular development from E6.5 to E9.5. Using this integrated atlas of mouse cardiovascular development, we also identified an Etv2 downstream target, *Ebf1*, as an important transcription factor for mouse endothelial development.

## Results

### Overview of single cell neighborhood component analysis

Our integration strategy aimed to correct the batch effects of temporal scRNA-seq data from multiple data sources while preserving the global developmental trajectory. Similar to the mutual nearest neighbors (MNN) based batch correction algorithms[12], we hypothesized that the cells that were mutually close to each other from two batches in the input space should be more likely to be similar cell types. However, instead of using a predefined number of neighbors per cell, we examined the distance between a pair of cells from two batches, by comparing their distance to a background distribution of cell distances (the null model), and the identification of the neighbors that were statistically and mutually similar to each other (see Materials and Methods and Fig S1A). Therefore, the neighbors per cell were automatically determined as the *context likelihood neighbors (CLN)*. It is known that for temporal scRNA-seq data, the time indices (i.e. the time stamp associated with each cell) is usually strongly associated with the gene expression pattern, leading to the difficulties of revealing the global trajectories and separating subpopulation of cells within each time point[10]. To account for the confounding effects of the time indices, the context likelihood was evaluated for the cells within each time point. Using synthetic temporal scRNA-seq data, we quantitatively compared the performance of capturing the true cell pairs from the same lineage, between CLN and MNN. Throughout this study, we utilized two types of synthetic temporal scRNA-seq data: *balanced* and *imbalanced* (see Materials and Methods). For the balanced data, each batch included the cells from all time points and all lineages. However, in a more realistic scenario, each data batch usually only covered partial time points and/or only included a subset of cell lineages. We referred to this type of data as the *imbalanced* temporal scRNA-seq data. We found that on both balanced and imbalanced synthetic scRNA-seq data, the CLN had significantly better performance for the identification of the same lineage of cells from different batches than by matching the MNN cells from different batches, as measured by the receiver operating characteristic (ROC) curves (Fig S1B-S1E).

Next, based on the cell-cell context likelihood from each time point, we estimated a batch-specific linear transformation **A** of the input space, [i.e. the gene expression matrix **X**, so that in the transformed space **AX** (i.e. the low dimensional space)], that the cells with high context likelihood between each other would be as close as possible (Fig 1A). We used a differentiable cost function based on stochastic neighbor assignment in the transformed space, similar to the neighborhood component analysis (NCA) [23]. It should be noted that although the CLN was evaluated for an individual time point, a single linear transformation applies to the cells from all time points. Thus, unlike our previous work on visualizing temporal scRNA-seq data[10], the cells from different time points will not be confined to the same transformed (low) dimensional space.

**Fig 1.**
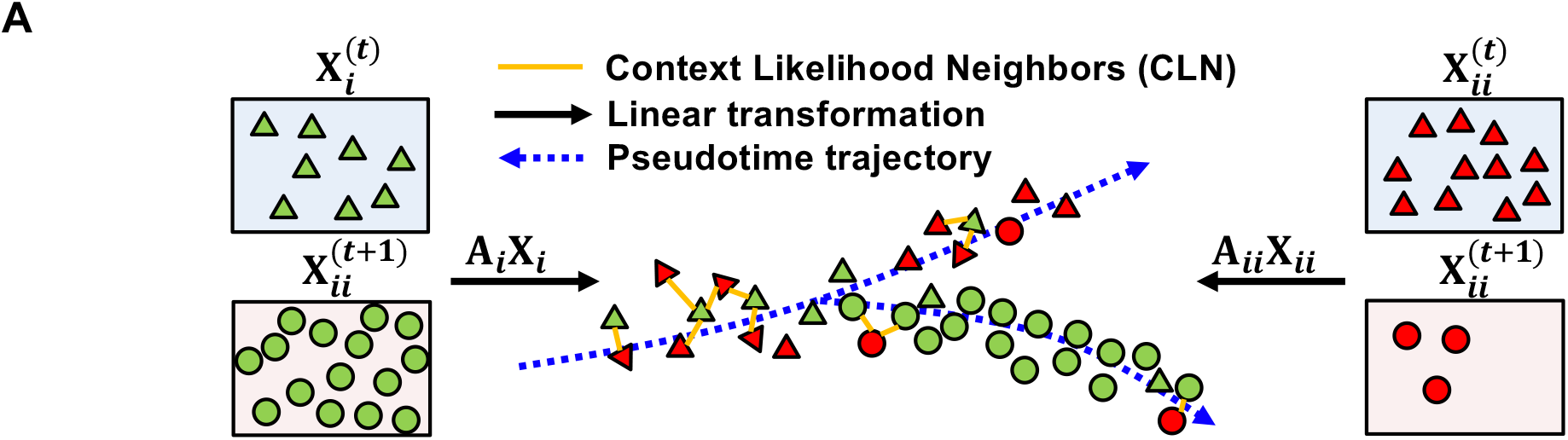
The overview of single cell neighborhood component analysis (scNCA). **(A)** scNCA first identified similar cell pairs from different batches at a specific time point using context likelihood as the context likelihood neighbors (CLNs). Then, scNCA learned a batch-specific linear transformation **A** of the input gene expression matrix **X**, so that in the transformed low dimensional space **AX**, the cells with high context likelihood between each other would be as close as possible.

### scNCA integrates imbalanced temporal scRNA-seq data

We first tested the performance of batch correction of scNCA on both balanced and imbalanced synthetic temporal scRNA-seq data with three lineages, three batches and five time points (Fig S2C and Fig 2C). A reversed graph embedding algorithm called DDRTree was used to visualize the single cell trajectories[24]. It has been shown that DDRTree excels at revealing the structure of complex trajectories with multiple branches from temporal scRNA-seq data. We utilized three different criteria to quantitatively compare the reconstructed lineage trajectories from uncorrected data, as well as data corrected by scNCA, along with two existing tools, scran’s mnnCorrect and Seura alignment[12,13]. These three criteria focused on three different aspects of batch correction: (1) how well cells from three batches mix together, (2) how well the global trajectories of three lineages were revealed, and (3) how well the cells from three lineages were locally grouped together. First, we devised a resampling-based statistical test for batch effects in temporal scRNA-seq data, tsBET (Fig S3). tsBET not only compares the local and global batch label distribution like the *k*-nearest neighbor batch effect test (kBET)[11], but also takes into account the confounding effects of time labels in the imbalanced temporal scRNA-seq data. Second, to investigate the global trajectories, we used *k*-means to cluster the cells on the DDRTree view (i.e. the 2D space produced by DDRTree algorithm) into three groups and compared the clusters with the known lineage labels using Adjusted Rand Index (ARI)[25]. Third, to examine the local distribution, we trained a multi-class linear support vector machine (SVM) using 2D coordinates of the DDRTree view as the feature, and the known lineage labels as the class labels, and computed the 10-fold cross validation accuracy[25]. Thus, high accuracy indicated the cells from the same lineage located together on the DDRTree view. For both balanced and imbalanced synthetic scRNA-seq data, we found that scNCA had significantly better performance of batch correction than mnnCorrect and Seurat alignment, on mixing cells from different batches, revealing the global trajectories, and bringing together the cells from the same lineage (Fig S2 for balanced data and Fig 2 for imbalanced data).

**Fig 2.**
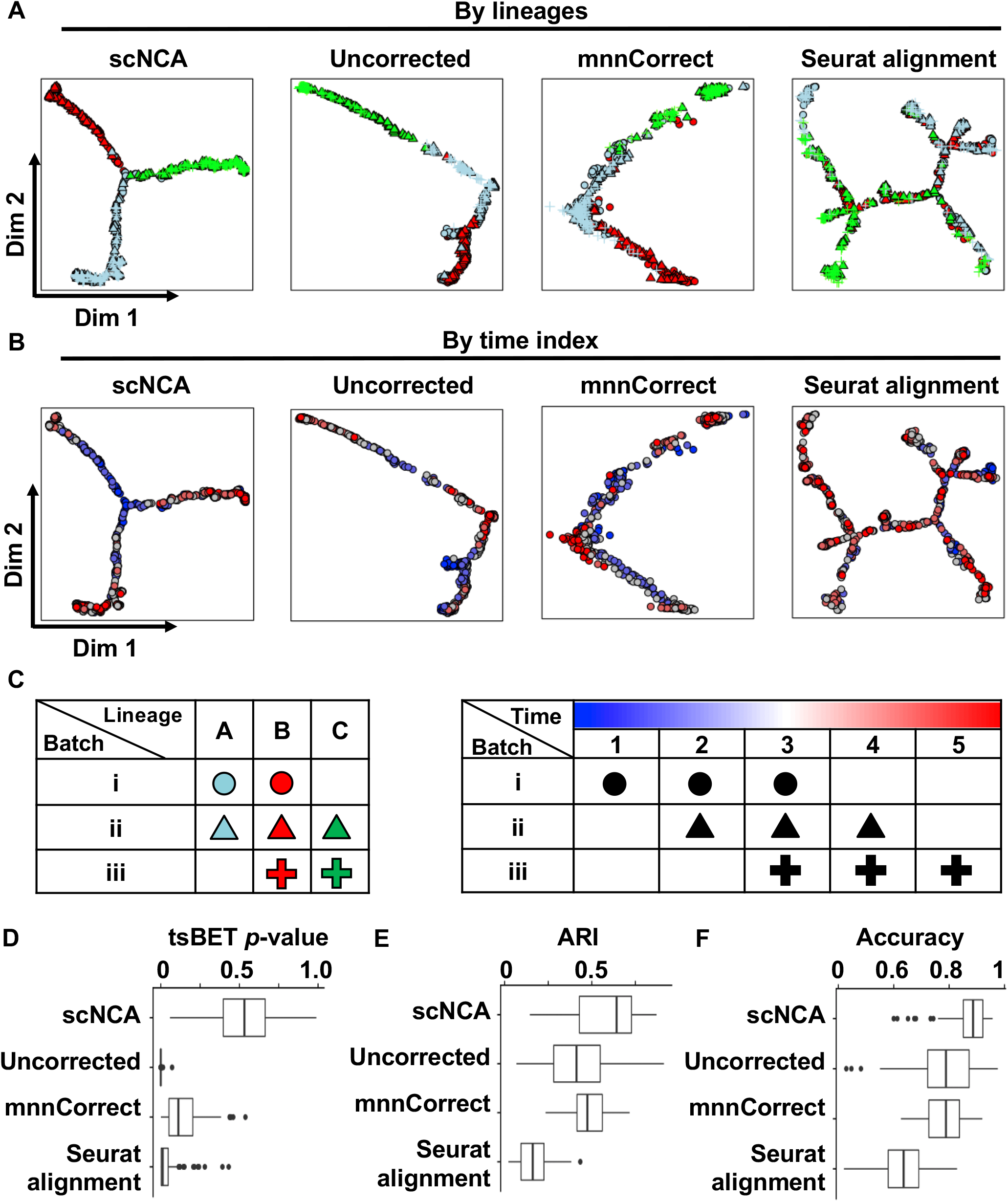
scNCA integrated the synthetic imbalanced temporal scRNA-seq data and preserved the lineage trajectories. **(A-B)** The two dimensional DDRTree view visualized the data integrated by scNCA, mnnCorrect and Seurat alignment, as well as the uncorrected data. In panel **(A)**, the color of the points indicates the lineages (light blue: lineage A; red: lineage B; green: lineage C). The shape of the points indicates the batches (circle: batch i; triangle: batch ii: cross: batch iii). In panel **(B)**, the color of the points indicates the associated time indices, from early time point (blue), to late time point (red). **(C)** In the synthetic imbalanced temporal scRNA-seq data, each batch includes only a subset of lineages, and/or covers partial time periods. **(D-F)** The performance of the batch correction is compared by **(D)** testing the significance of a local mixture of cells from different batches (tsBET *p*-values), **(E)** the adjusted Rand Index between *k*-means clustering results of cells on the DDRTree view and the known lineage labels, and **(F)** the 10-fold cross validation accuracy of a multi-class linear support vector machine (SVM) using 2D coordinates of the DDRTree view as the features and the known lineage labels as the class labels. Each comparison was repeated 100 times.

Having demonstrated the superior performance of scNCA using synthetic datasets, we continued to investigate its performance using two real imbalanced temporal scRNA-seq datasets. We first used scNCA to integrate two scRNA-seq datasets of mouse early embryonic development[1,2]. Although tsBET showed that there was no significant batch effects in each of the DDRTree view, we found that the scRNA-seq dataset integrated by scNCA formed a clear bifurcation into two distinct lineages, trophectoderm (TE) and inner cell mass (ICM), as evidenced by the Sox2/Cdx2 ratio of blastocyst cells (Fig S4). In the uncorrected and data corrected by mnnCorrect, the TE and ICM lineages were not separated. While in the data corrected by Seurat alignment, the TE and ICM did separate, the blastocyst cells from Deng et al. dataset (red dots in Fig S4B) were mistakenly mixed with the earlier stage cells from the Goolam et al. dataset, at the tips of TE and ICM trajectories.

Next, we used scNCA to integrate 3,387 single cells from six published temporal scRNA-seq datasets focused on mouse cardiovascular development from the epiblast (E6.5) to the four-chambered heart (E9.5) (Fig 3C). These datasets included the single cells from the whole epiblast[3–5], Mesp1^+^ cardiovascular progenitors[4], Flk1^+^ mesodermal progenitors[5], Etv2^+^ hemato-endothelial progenitors[6], as well as two scRNA-seq datasets obtained from distinct cardiac regions at E8.5 and E9.5[7,26]. We also added 264 Nkx2-5-EYFP^+^ single cells from the E7.75 and E8.5 mouse embryo. Previous analysis showed that Nkx2-5 was expressed in multipotent progenitors in the mouse embryo, and targeted disruption of Nkx2-5, in the mouse, resulted in perturbed heart morphogenesis and perturbed endothelial and hematopoietic development at approximately E9.5[27]. Thus, the Nkx2-5-EYFP^+^ scRNA-seq datasets from E7.75 and E8.5 bridged the gap between early cardiovascular progenitors represented by Mesp1^+^, Flk1^+^ and Etv2^+^ cells, and the late whole heart scRNA-seq dataset from E9.5.

**Fig 3.**
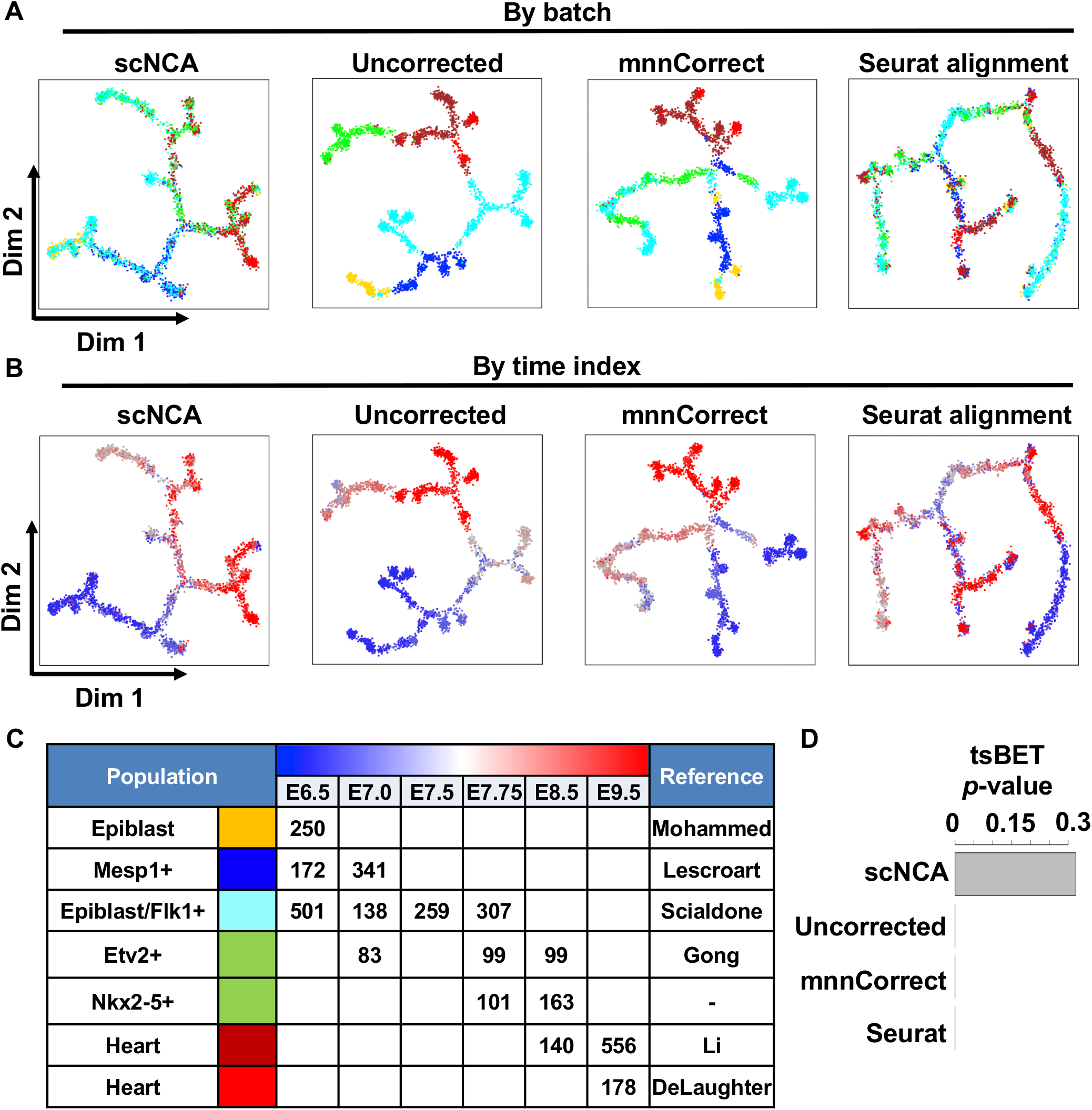
scNCA integrated 3,387 single cells from seven heterogenous temporal scRNA-seq datasets during mouse cardiovascular development from E6.5 to E9.5. **(A-B)** The two dimensional DDRTree view visualized the data integrated by scNCA, mnnCorrect and Seurat alignment, as well as the uncorrected data. In panel **(A)**, the color of the points indicates the data sources, while in panel **(B)**, the color of the points indicates the associated developmental stages, from an early time point E6.5 (blue), to a relatively late time point E9.5 (red). **(C)** The table shows the distribution of single cells across seven different studies and six different developmental stages. **(D)** The tsBET for the distributions of the integrated cells on DDRTree view shows that there is no significant batch effects of the data corrected by scNCA, while there are significant batch effects of the data corrected by mnnCorrect, Seurat alignment, as well as the uncorrected data.

Visual inspection of the DDRTree view produced by the data corrected by scNCA showed not only that the cells from different studies were well mixed (Fig 3A) but also a clear gradient of cells from early (blue cells in Fig 3B) to late stages (red cells in Fig 3B). The DDRTree view produced by uncorrected data and the data corrected by mnnCorrect had clear clustering of cells from the same studies (batches) (Fig 3A). Similar to scNCA, Seurat alignment also mixed the cells well, but the gradient of the developmental stages was lost (Fig 3B). Indeed, the tsBET analysis showed that there was no significant batch effects in the data integrated by scNCA (tsBET *p*-value=0.312), while the batch effects in the uncorrected data, and the data corrected using mnnCorrect and Seurat alignment were significant (tsBET p-value < 1e-10) (Fig 3D).

Close examination of the data integrated by scNCA, we found that the cardiac muscle marker, Tnnt2, was highly expressed on the right side of the DDRTree view (Fig 4A), along with left ventricle marker, Myl2, and the outflow tract marker, Isl1 (Fig S5A and S5B). In contrast, the Tnnt2-high populations in the data corrected by Seurat alignment were distributed at distant branches, suggesting the failure of merging related cardiac cells (Fig S5C). Previous studies showed that the Nkx2-5^+^ cells at E7.75 give rise to endothelial and smooth muscle cells in the E9.5 mouse heart[27]. Thus, we examined the Nkx2-5-EYFP^+^ cells at E7.75 and the cells of whole heart tissues at E9.5 on the DDRTree view (Fig 4B and S5D). We hypothesized that overall distance between E7.75 Nkx2-5^+^ (progenitor) cells and E9.5 (differentiated) cardiac cells should be relatively close to each other on well-integrated data. Indeed, the overall distance between each Nkx2-5^+^ cell and its nearest E9.5 cardiac cell on the DDRTree view produced by scNCA was significantly smaller than that produced by uncorrected, and data corrected by mnnCorrect and Seurat alignment (Fig 4C). Taken together, these results based on visual inspection, statistical test of batch effects (tsBET) and the biological facts consistently suggested that scNCA demonstrated superior performance on the integration of the imbalanced temporal scRNA-seq datasets focused on early cardiovascular development in the mouse.

**Fig 4.**
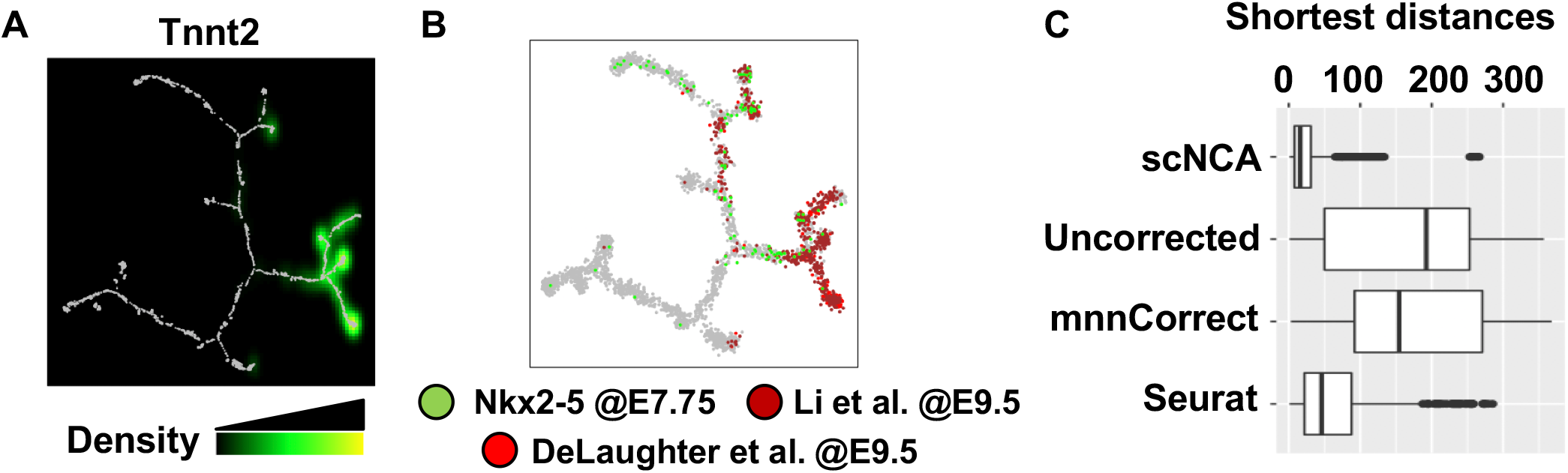
scNCA corrected the batch effects of Nkx2-5+ cell and E9.5 cardiac cell on integrated data. **(A)** The Tnnt2 expression level of the data integrated by scNCA indicates that the cells from cardiac lineages are enriched at the right side of the DDRTree view. The expression level of Tnnt2 is shown as the density of normalized read counts of Tnnt2 (black: low expression level; green: medium expression level; yellow: high expression level). **(B)** The DDRTree view indicates the Nkx2-5-EYFP+ cells from E7.75 (green points), and whole heart cells from E9.5 (brown and red points). **(C)** The boxplot compares the distribution of the distance from E7.75 Nkx2-5-EYFP+ cells to their nearest E9.5 whole heart cells on the DDRTree view of the data integrated by scNCA, mnnCorrect and Seurat alignment, as well as the uncorrected data. The distances between two groups of cells on the data integrated by scNCA are significantly less than other three methods (Wilcoxon signed rank test *p*-value < 0.001).

### The Etv2 downstream target gene, *Ebf1*, promotes endothelial development

We examined the expression of Etv2-EYFP^+^ cells and the known hemato-endothelial markers and we identified branches of endothelial and hematopoietic lineages on the integrated cardiovascular developmental datasets (Fig 5A and Fig S6). We further found that among 78 transcriptional factors that have Etv2 ChIP-seq binding sites within a 5-kb region surrounding their transcriptional start sites, the Early B-Cell Factor 1 (Ebf1) had one of the highest expression levels in the endothelial branch compared with the hematopoietic branch (Fig 5B and 5C)[28]. Ebf1 has been previously shown to control B cell differentiation and expressed in the cardiopharyngeal mesoderm[29,30]. However, the functional role of Ebf1 in early mesodermal development remains unclear. First, we observed that the *Ebf1* gene harbored an evolutionary conserved Etv2 binding motif in the upstream region (Fig S7A). Using qPCR analysis and Doxycycline-inducible HA-tagged Etv2 ES/EB system, we observed increased expression of Ebf1 following the induction of Etv2 relative to uninduced EBs (Fig 5D). Next, our transcriptional assays using the 0.5kb *Ebf1* promoter-reporter construct revealed that Etv2 potently transactivated the *Ebf1* promoter in a dose-dependent fashion. Mutagenesis of the Etv2 binding motif resulted in abolishment of the transcriptional activity (Fig 5E). The gel-shift assays further confirmed that Etv2 could bind to the *Ebf1* promoter containing the Etv2 recognition sequence (Fig 5F). To visualize whether Ebf1 was expressed in the mesodermal progenitors, we sorted Flk1^-^ and Flk1^+^ cells from differentiating EBs and performed qPCR analysis for Ebf1 transcripts. Our analysis revealed that Ebf1 was significantly enriched in the Flk^+^ cells as compared to Flk1^-^ cells (Fig S7B and S7C). To further investigate the function of Ebf1 in endothelial development, we utilized the Dox-inducible Ebf1 ES/EB system and FACS analysis for endothelial lineages. Addition of Dox resulted in ~1.5 fold increase in the percentage of Ebf1^+^ cells (Fig S7D). Finally, our FACS analysis revealed that the over-expression of Ebf1 resulted in a significant increase of the endothelial program (CD31^+^/Tie2^+^ and CD31^+^/Flk1^+^ populations) in +Dox as compared to −Dox treated EBs (Fig 5G-5I; Fig S7E-S7G). Thus, by using integrated scRNA-seq datasets from early cardiovascular development in the mouse, we successfully identified and experimentally verified that Ebf1, was a direct downstream target of Etv2, which was highly expressed in endothelial lineages and promoted endothelial development.

**Fig 5.**
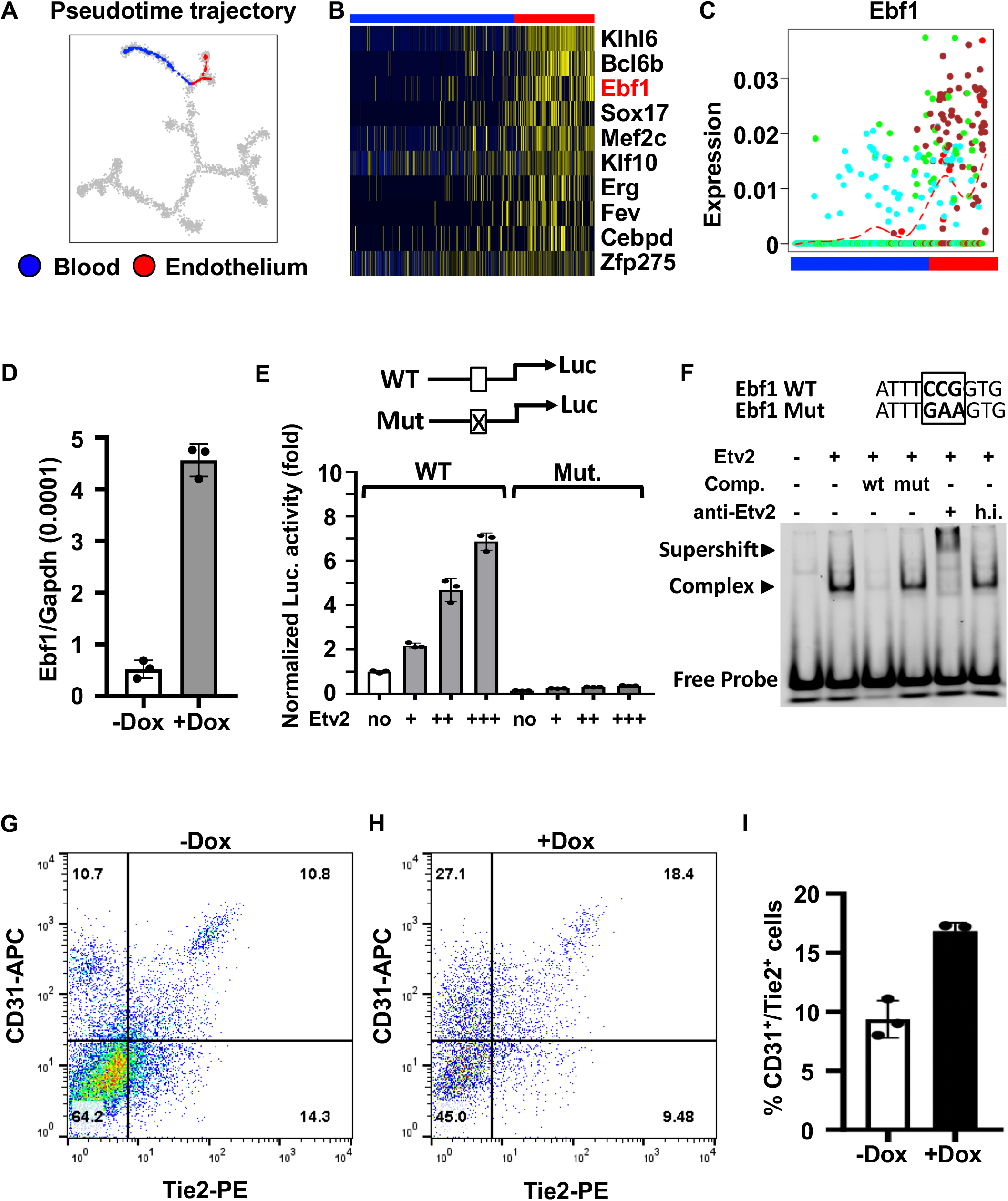
The Etv2 downstream target gene, *Ebf1*, promotes endothelial development. **(A)** The DDRTree view shows the blood (blue) and endothelial (red) lineages of the data integrated by scNCA. **(B)** The Early B-Cell Factor 1 (Ebf1) has one of the highest expression levels in the endothelial branch compared with the hematopoietic branch. The color in the heatmap indicates the standardized gene expression levels (blue: low expression levels; yellow: high expression levels). **(C)** The normalized expression levels of Ebf1 along the blood (blue) and endothelial (red) branches. The y-axis shows the cosine normalized expression levels. **(D)** qPCR analysis shows significantly increased expression of *Ebf1* following the induction of Etv2 relative to uninduced EBs. **(E)** Luciferase reporter assays using the *Ebf1* promoter (0.5 kb) harboring wild-type (WT) or mutant (mut) Etv2 binding site shows that Etv2 potently transactivates the *Ebf1* promoter in a dose-dependent fashion. **(F)** Gel-shift assay shows that Etv2 binds to the Ets binding site in the *Ebf1* promoter region. Moreover, the Etv2 binding to this binding site can be competed and super-shifted. **(G-H)** FACS analysis and **(I)** quantification of endothelial lineages of endothelial markers (Tie2 and CD31) in −Dox and +Dox conditions indicate that over-expression of Ebf1 results in significant increase of the endothelial program. Error bars indicate SEM.

## Discussion

As single cell RNA-seq techniques have been widely used to study focused stages or subpopulations of cells of complex biological process such as cardiovascular development, the integration of these separate datasets will produce a broader and more comprehensive view of the underlying molecular dynamics that could not be possibly revealed using the data from much shorter time periods. Here, we provide evidence that scNCA integrates heterogenous synthetic and real temporal scRNA-seq datasets from multiple sources with strong batch effects. scNCA not only successfully corrected the batch effects, but also preserved the global structure of gene expression. The time dependent context likelihood that scNCA utilized for matching similar cells from different batches successfully eliminated the confounding effects of time indices and resulted in significantly better performance of matching cells from the same lineages. We believe that the same principal can be potentially applied to address other known confounding factors, such as sequencing platforms, other than time indices.

Although non-linear transformation such as deep generative models have been applied to transform the scRNA-seq data into low dimensional space[31,32], scNCA employed a linear transformation strategy and resulted in superior performance for the correction of batch effects. Nevertheless, more complicated non-linear transformation could be explored under the framework of neighborhood component analysis.

In summary, we presented scNCA as a novel tool to correct the batch effect of temporal scRNA-seq. We used scNCA to integrate 3,387 single cells from seven heterogenous temporal scRNA-seq datasets of mouse early cardiovascular development, and identified an Etv2 downstream target, *Ebf1*, as an important transcription factor for mouse endothelial development. We provide the R/TensorFlow implementation of scNCA at https://github.com/gongx030/scNCA. The integrated mouse early cardiovascular development data can be explored at https://heartmap.umn.edu/scNCA.

## Materials and Methods

### Notations

We first defined the main notations used for representing the temporal the scRNA-seq data. For a temporal scRNA-seq dataset with *N* genes, *B* batches, *T* time points and *M* cells, let 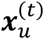 be the gene expression matrix of cell *u* at time *t* ∈ {1, …, *T*}. The batch label of cell *u* at time *t* is represented by 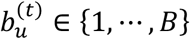. The number of cells from time *t* is 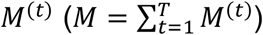. We use **X***_i_* to denote the gene expression matrix of cells from batch *i* ∈ {1, …, *B*}.

### Simulating balanced and imbalanced temporal scRNA-seq data

We first defined the number of genes *N* = 2000, number of lineages *H* = 3(lineage A, B, and C), the number of batches *B* = 3 (batch *i*, *ii*, and *iii*), and the number of time points *T* = 5 (time index 1, 2, 3, 4 and 5). For the balanced temporal scRNA-seq data where each batch had the cells from all the lineages, we assumed each batch had 100 cells from each of the lineages, resulting in total number of cells *M* = 900 (Fig S2C). For the imbalanced data where each batch only had a subset of lineages, we assumed that batch *i* had 100 cells from lineage A and B, respectively, batch *ii* had 100 cells each of three lineages, and batch *iii* had 100 cells from lineage B and C, resulting in total number of cells *M* = 700 (Fig 2C).

The simulation of temporal scRNA-seq datasets followed three basic assumptions: (1) the developmental trajectories could be represented within a *K*-dimensional biological subspace. (2) We assumed that all *H* lineages arose from the same progenitors and differentiated toward different directions. Thus, to simulate the developmental trajectory of any lineage *h* ∈ {1, …, *H*}, we used the origin of the *K*-dimensional space as the starting point, we randomly selected another point in the *K*-dimensional space as the terminal point, and drew a *lineage segment* (a line segment) that connected the origin and the terminal point. This lineage segment therefore represented the developmental trajectory of lineage *h* on the *K*-dimensional biological subspace. (3) The developmental speed was constant. Thus, the each lineage segment was evenly split into *T* parts, where each part was associate with the time index from 1 to *T*. To simulate the cells of lineage *h* from time *t* ∈ {1, …, *T*}, we randomly drew points from the lineage segments that corresponded to lineage *h* from time *t* on the *K-*dimensional space. Note that for balanced data, the points were drawn from the entire lineage segments, while for the imbalanced data, the points could only be drawn from the lineage segments covered by the corresponding time period. Then to simulate high-dimensional gene expression, the *K*-dimensional points were projected to *N*-dimensional space by a randomly Gaussian matrix. Similar to Haghverdi et al., the batch effects were incorporated by generating a Gaussian random vector for each batch and adding it to the gene expression matrix[12].

### The context likelihood

We used the context likelihood to estimate the likelihood of the Euclidean distance *d_uv_* for a particular pair of cells *u* and *v* from batch *b_u_* and *b_v_*, respectively, by comparing the background distribution of Euclidean distance (the null model). The background distribution was constructed from two sets of distance values: {*d_u_*}, the set of the distance values from cell *u* to all cells *not* from batch *b_u_*, and {*d_v_*}, the set of the distance values from cell *v* to all cells *not* from batch *b_v_*. The background distance distribution *d_uv_* was approximated as a joint normal distribution with *d_u_* and *d_v_* as the independent variables. Thus, the final form of the context likelihood between cells *u* and *v* is 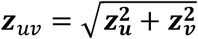, where ***z**_u_* and ***z**_v_* are the *z*-scores of *d_uv_* from the marginal distributions (Fig S1A). By default, we used a 20% quantile (*p* = 0.2) of all elements in ***Z*** (*c*_0_) as the threshold for neighboring cells. Finally, we defined a *context likelihood neighbor*(CLN) matrix **W** for all pairs of cells such that:

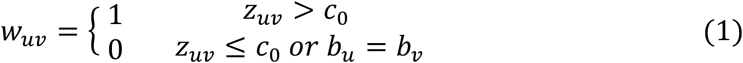

Note that the CLNs were computed for all cells from each time point *t* to account for the confounding effects of time indices and the CLNs for time *t* was denoted as **W**^(*t*)^. The scheme above was similar to the context likelihood of relatedness (CLR) algorithm for discovering gene regulatory networks[33].

### Neighborhood component analysis for single cell RNA-seq (scNCA)

For each batch *i* ∈ {1, …, *B*}, we learnt a *K × N* linear transformation ***A**_i_* of the input space ***X**_i_*, so that in the transformed space ***A**_i_**X**_i_*, neighboring cells with high context likelihood would be as close as possible (Fig 1A). The *K* was the number of low dimensions used for scNCA and we used *K* = 5 throughout this study. Specifically, we defined a probability 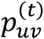 as the probability of cell *u* as the neighbor of cell *v* in the transformed space at time *t*, and 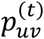 could be computed as a softmax over Euclidean distances in the transformed space:

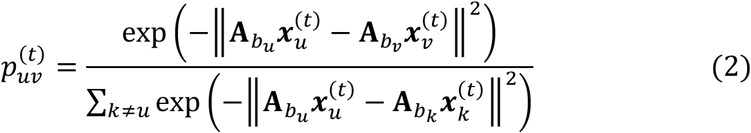

Then we computed the probability 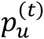 that cell *u* would be correctly grouped with the its CLNs at time *t*:

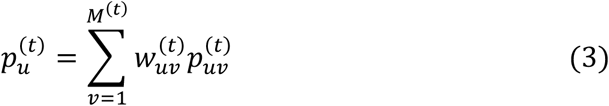

And finally, the objective we maximized was the expected number of CLNs in the transformed space over all *T* time points:

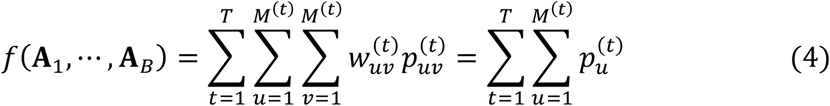

We used the adaptive gradient algorithm AdaGrad for optimizing the objective function with a default learning rate of 0.01[34]. The optimization was performed by using TensorFlow 1.12.

### Testing for the batch effects in temporal scRNA-seq data (tsBET)

The real temporal scRNA-seq data usually showed an imbalance of between the distribution of time indices and batch labels associated with each cell. To test the batch effects in the transformed low dimensional space (e.g. the DDRTree view), the local distribution of time indices and batch labels surrounding each cell were examined at the local branching structure. First, we defined *g*(*u, k*) as the set of all *pairs* of neighboring cells in the *k* -local branching structure surrounding cell *u*. For the DDRTree view, *k* -local branching structure was *k* nearest cells to cell *u* on the global minimum spanning tree[24]. We used a batch score *s_u_* to measure the local batch effects surrounding cell *u*, whereby:

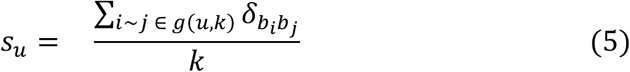

where *δ_ij_* was the Kronecker delta function, equaling to 1 if *i* = *j*, and 0 otherwise. To account for the global imbalanced distribution between time indices and batch labels, for the *k* -local branching structure surrounding cell *u*, a null distribution of batch score, 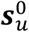, was computed by using the batch labels that were randomly generated by the time indices based on the global time/batch distributions. This random sampling was repeated for 100 times for *k* -local branching structure of each cell *u*. Thus, the significance of batch effect of the local branching structure could be computed as:

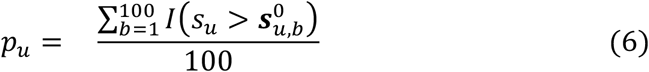

where *I*(*x*) was the indicator function, equaling 1 if the *x* is true, and 0 otherwise. The significance of global batch effects was computed by aggregating the significance of local batch effects:

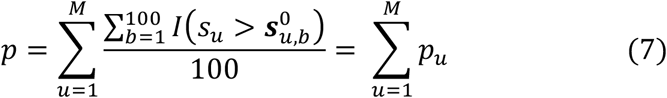

The choice of the size *k* of the local branching structure mainly depended on the size of the dataset. In this study, we used *k* = 100 for the synthetic data (900 and 700 cells for balanced and imbalanced data, respectively) and the cardiovascular development data (3,387 cells). We used *k* = 20 for the mouse preimplantation embryonic development data, which had a relatively smaller size of 354 cells.

### Processing of scRNA-seq data

The raw reads of the public datasets used in this study were downloaded from NCBI SRA (Deng et al.: GSE45719[1]; Mohammed et al.: GSE100597[3]; Lescroart et al.: GSE100471[4]; Gong et al.: PRJNA350294[6]; Li et al.: GSE76118[35]), EBI ArrayExpress (Goolam et al: E-MTAB-3321[2]; Scialdone et al: E-MTAB-4079[5];) and GNomEx databases (DeLaughter: 272R, 274R, 275-292R, 439R, and 440R[7]). The raw read counts for each gene were obtained with TopHat (v2.0.13) and HTSeq (v0.6.0) with default parameters[36,37]. The raw read counts were first normalized by the devolution-based size factors, as implemented by the function *computeSumFactors* in the *scran* package[38]. Then, we used the *trendVar* function in the *scran* package to fit a mean-variance trend model using endogenous genes. The highly variable genes (HVGs) with a false discovery rate of 5% or less were used for the integration analysis. There were 2,617 and 4,904 HVGs for the mouse preimplantation embryonic data, and the mouse cardiovascular development data, respectively. For the analysis of uncorrected data, scNCA or mnnCorrect (*k*=20), the remaining data were then scaled through a cosine normalization, due to its robustness to technical differences in sequencing depth and capture efficiency between batches[12]. For Seurat alignment, we used the raw read counts of HVGs as the input, followed by the internal normalization and scaling (*NormalizeData* and *ScaleData* implemented in the Seurat package). A CCA (canonical component analysis) was performed on the resulting data (*RunCCA* with *num.cc=5*), followed by the subspace alignment (*AlignSubspace*)[13].

### Isolation of Nkx2-5-EYFP^+^ single cells

Nkx2-5-EYFP embryos were harvested from time mated females at embryonic day (E)7.75 or E8.25 and screened using microscopy for EYFP expression[27]. Embryos were divided into EYFP positive and negative pools for dissociation with TrypLE Express (Gibco by Life Technologies). After dissociation, cells were diluted with 10% FBS in DMEM and pelleted at 1000G. Cells were resuspended in PBS containing 2% serum and 0.1%propidium iodide (PI). EYFP negative embryos were used as a gating control sample. PI negative, EYFP positive cells were sorted by FACS using a MoFlo XDP (Beckman Coulter) into DMEM plus 10% FBS. FACS sorted cells were resuspended at 500 cells/μl before loading onto a Fluidigm 10-17μm integrated fluidics circuit (IFC) for capture, viability screening, lysis, and library amplification on a C1 Single-Cell Auto Prep System (Fluidigm).

### Single cell RNA-seq of Nkx2-5-EYFP^+^ cells

All libraries were sequenced using 75-bp paired end sequencing on MiSeq (Illuminia). The cells with less than 100K paired reads were removed, resulting in 315 cells for analysis. The raw read counts for each gene were obtained with TopHat (v2.0.13) and HTSeq (v0.6.0) with default parameters[36,37]. The median mapping rate was 89.2%. The raw read counts were normalized by the devolution-based size factors, as implemented by the function *computeSumFactors* in the *scran* package[38].

### Electrophoretic mobility shift assay (EMSA)

pcDNA3.1-Etv2-HA or empty pcDNA3.1(+)-HA vectors were expressed using the TNT Quick Coupled Transcription/Translation System (Promega, Madison, WI) according to the manufacturer’s protocol. Oligo DNA corresponding to the wild-type *Ebf1* promoter sequence or an oligo DNA sequence having a CCG to GAA mutation in the putative Etv2 binding site were synthesized with and without the IRDye^®^ 700 fluorophore (Integrated DNA Technologies, Coralville, IA). Ebf1 WT top labeled: IRD700-AGTCCGGATTTCCGGTGGCGTTCT; Ebf1 WT top: AGTCCGGATTTCCGGTGGCGTTCT; Ebf1 WT bottom: AGAACGCCACCGGAAATCCGGACT; Ebf1 mut top: AGTCCGGATTTGAAGTGGCGTTCT; Ebf1 mut bottom: AGAACGCCACTTCAAATCCGGACT. Complimentary WT or mutant oligos were annealed to generate the labeled probe and unlabeled competitor DNA. In vitro synthesized protein (1μL) was incubated with 250 ng of poly dI-dC in binding buffer (50 mM Tris pH 7.6, 80 mM NaCl, 8 % glycerol) at room temperature for 10 minutes. For the antibody treatment, pre-binding was performed in the presence of active or heat-inactivated anti-human Etv2 antibody (ER71 (N-15), catalog #sc-164278; Santa Cruz Biotechnology, Inc., Dallas, TX). IRDye^®^ 700-labelled probe (100 fmol) was then added and the binding reaction proceeded at room temperature for 15 minutes. DNA-protein complexes were resolved on a 6% non-denaturing polyacrylamide gel in 0.5x TBE (40 mM Tris pH 8.3, 45 mM boric acid, and 1 mM EDTA) at room temperature. Fluorescence was detected using an Odyssey CLx imager (LI-COR Biosciences, Lincoln, NE).

### Luciferase assays

The *Ebf1* promoter region (0.5kb) and its mutant harboring the Ets binding site (EBS) was amplified by PCR and subcloned into the pGL3 vector to generate the *Ebf1* promoter-reporter. Cos-7 cells were cultured in Dulbecco’s modified Eagle’s medium complete medium supplemented with 10% FBS and 1X penicillin/streptomycin (ThermoFisher Scientific). Cells were trypsined using 0.25% trypsin and 1X10e5 cells per well were plated in a 12 well plate and cotransfected using Lipofectamine 3000 (Life Technologies) with increasing amounts of Etv2 plasmid and wild-type (WT) or mutant (mut) promoter-reporter constructs. 10 ng of pRL-CMV (Promega) was used as an internal control. Following transfection (36 h), cells were harvested and luciferase activity was analyzed with Dual Luciferase Stop-Glo System (Promega) and normalized with the Renilla luciferase.

### Embryonic stem (ES) cell culture and Embryoid body (EB) differentiation

Doxycycline-mediated Etv2- and Ebf-overexpressing mouse embryonic stem (mES) cells were generated using an inducible cassette exchange strategy. mES cells were differentiated into embryoid bodies (EBs) using mesodermal differentiation media containing, 15% FBS (Foundation ES Cell serum), 1× penicillin/streptomycin, 1× GlutaMAX (Gibco), 50 μg/ml Fe-saturated transferrin, 450 mM monothioglycerol, 50 μg/ml ascorbic acid in IMDM (Invitrogen) using the shaking method. Doxycycline was added between D2-D4 after initiation of differentiation and harvested for the respective experiments.

### Fluorescence-activated cell sorting (FACS) and analysis

Differentiating embryoid bodies were dissociated using 0.25% trypsin and single cell suspension were used for FACS staining. The antibodies used for FACS include: Ebf1-PE (BD Biosciences, 565494), Flk1-PECy7 (BD Biosciences, 561259), Tie2-PE (eBiosciences, 12-5987-82), and CD31-APC (eBiosciences, 17-0311-82). Stained cells were sorted and analyzed using a FACSAria machine and the data were processed using FlowJo software version 10.3.

### RNA isolation and qPCR

Flk1-FACS sorted cells, or the whole EB cells were lysed in RLT lysis buffer and total RNA was isolated using RNeasy kit (Qiagen) according to the manufacturer’s protocol. cDNA was synthesized using SuperScript IV VILO kit (Thermo Fisher Scientific) and quantitative PCR (qPCR) assay was performed with ABI Taqman probe sets.

### Availability of data and materials

The single cell RNA-seq data that support the findings of this study have been deposited in NCBI Sequence Read Archive (SRA) database with the project accession number PRJNA509274. The *scNCA* software was freely available at https://github.com/gongx030/scNCA. The All other relevant data are available from the authors. The scNCA-integrated mouse early cardiovascular development data can be explored at https://z.umn.edu/scNCA.

## Acknowledgments

Funding support was obtained from the National Institutes of Health (U01HL100407 to D.J.G), the Department of Defense (GRANT11763537) and Regenerative Medicine of Minnesota. We acknowledge the support from the University of Minnesota Supercomputing Institute.

## Author Contributions

D.J.G. conceived the study. W.G. designed and implemented the computational approach, analyzed the data and drafted the manuscript. P.S. compiled the scRNA-seq for the analysis, designed http://heatmap.umn.edu. B.N.S., J.T. and S.D. performed the experiments. W.P. and D.J.G. supervised the study. All authors (W.G., B.N.S., P.S., S.D., J.T., S.C, M.K., M.G.G., D.Y., W.P., D.J.G.) read, edited and approved the final manuscript.

## Conflict of Interest Statement

The authors declare no competing interests.

