## Supplementary Information for "A novel algorithm for the collective integration of single cell RNA-seq during embryogenesis"

Fig S1

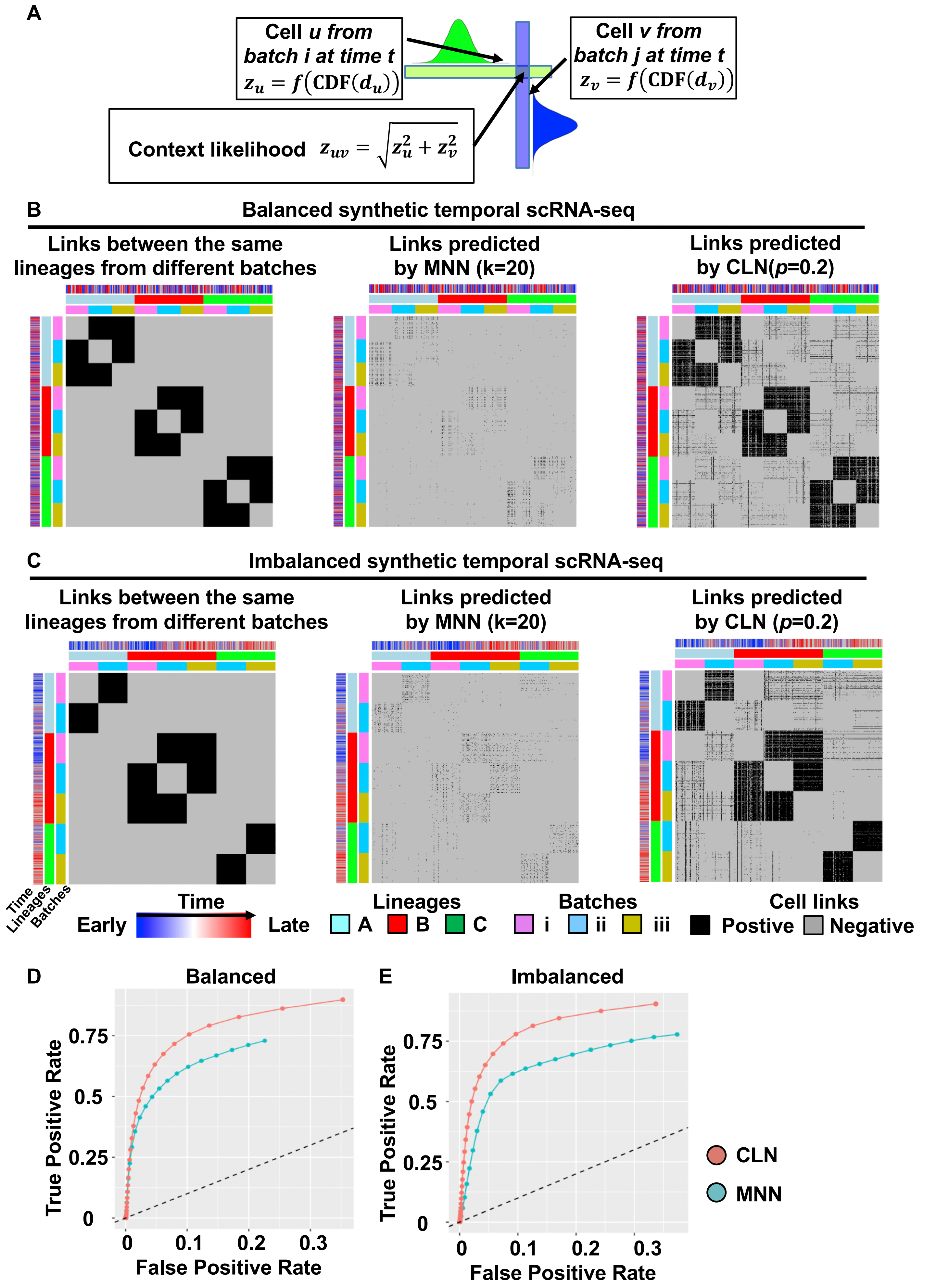

A
Cell u from batch i at time t
$z_u = f(\mathrm{CDF}(d_u))$
Cell v from batch j at time t
$z_v = f(\mathrm{CDF}(d_v))$
Context likelihood $z_{uv} = \sqrt{z_u^2 + z_v^2}$
B
Balanced synthetic temporal scRNA-seq
Links between the same lineages from different batches
Links predicted by MNN (k=20)
Links predicted by CLN(p=0.2)
C
Imbalanced synthetic temporal scRNA-seq
Links between the same lineages from different batches
Links predicted by MNN (k=20)
Links predicted by CLN (p=0.2)
Time
Lineages
Batches
Time
Early
Late
Lineages
A
B
C
Batches
i
ii
iii
Cell links
Postive
Negative
D
Balanced
E
Imbalanced
True Positive Rate
False Positive Rate
0
0.25
0.5
0.75
0.1
0.2
0.3
CLN
MNN

**Fig S1. The time dependent context likelihood captures the cells from the same lineages among different batches of temporal scRNA-seq data. (A)** The schematic illustrates how context likelihood is computed between cells from different batches. At a specific time point $t$, the $z$-score of all possible neighbors of cell $u$ from batch $i$ ($\boldsymbol{z}_u$) is computed according the distribution of Euclidean distance between cell $u$ and all cells outside of batch $i$, in the input space. The $z$-score of all possible neighbors of cell $v$ from batch $j$ ($\boldsymbol{z}_v$) is computed in the similar fashion. The context likelihood of cell $u$ and cell $v$ ($\boldsymbol{z}_{uv}$) is computed as the joint likelihood estimates in the form of $\boldsymbol{z}_{uv} = \sqrt{\boldsymbol{z}_{\boldsymbol{u}}^{\boldsymbol{2}} + \boldsymbol{z}_{\boldsymbol{v}}^{\boldsymbol{2}}}$. **(b-c)** The heatmaps show the known (simulated) links of the cells from the same lineages between different batches (left column), the links predicted by mutual nearest neighbor method (MNN) using the default number of neighbors $k = 20$ (middle column), and the links predicted by context likelihood using the default parameter $p = 0.2$ (right column), for the **(B)** balanced and **(C)** imbalanced synthetic data. In the main heatmaps, the black and gray colors indicate the positive and negative links, respectively. The row and column annotations indicate the time indices from early time point (blue), to late time point (red), lineages (light blue: lineage A; red: lineage B; green: lineage C), and the batches (pink: batch i; cyan: batch ii: yellow: batch iii). **(D-E)** The receiver operating characteristic (ROC) illustrates the false positive rate (FPR) (*x*-axis) and the true positive rate (TPR) (*y*-axis) of using context likelihood and MNN to recover the cells from the same lineage on **(D)** balanced and **(E)** imbalanced synthetic temporal scRNA-seq data. For MNN, the number of neighbors $k$ ranges from 10 to 200. For context likelihood, the parameter $p$ ranges from $10^{-5}$ to 1. For each configuration, the temporal scRNA-seq data are randomly simulated for 100 times and the mean FPR and TPR are shown.

Fig S2

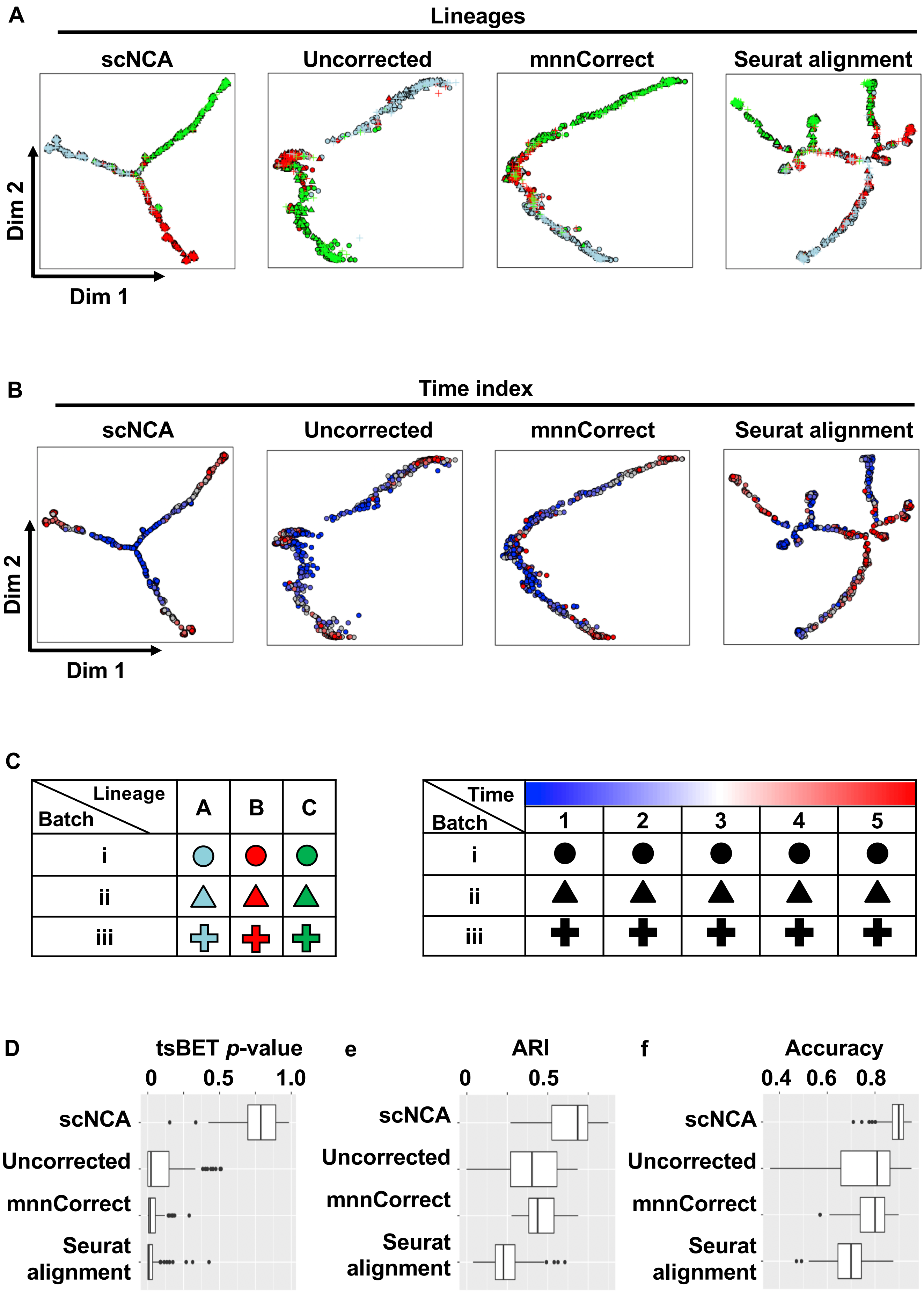

A
Lineages
scNCA
Uncorrected
mnnCorrect
Seurat alignment
Dim 2
Dim 1
B
Time index
scNCA
Uncorrected
mnnCorrect
Seurat alignment
Dim 2
Dim 1
C
Lineage
Batch
A
B
C
i
ii
iii
Time
Batch
1
2
3
4
5
i
ii
iii
D
tsBET p-value
0
0.5
1.0
scNCA
Uncorrected
mnnCorrect
Seurat alignment
e
ARI
0
0.5
scNCA
Uncorrected
mnnCorrect
Seurat alignment
f
Accuracy
0.4
0.6
0.8
scNCA
Uncorrected
mnnCorrect
Seurat alignment

**Fig S2. scNCA integrates the synthetic balanced temporal scRNA-seq data and preserves the lineage trajectories. (A-B)** The two dimensional DDRTree view visualizes the data integrated by scNCA, mnnCorrect and Seurat alignment, as well as the uncorrected data. In panel **(A)**, the color of the points indicates the lineages (light blue: lineage A; red: lineage B; green: lineage C). The shape of the points indicates the batches (circle: batch i; triangle: batch ii: cross: batch iii). In panel **(B)**, the color of the points indicates the associated time indices, from early time points (blue), to late time points (red). **(C)** In the synthetic balanced temporal scRNA-seq data, each batch covers the cells from all possible lineages and time points. **(D-G)** The performance of the batch correction is compared by **(E)** testing the significance of a local mixture of cells from different batches (tsBET *p*-values), **(F)** the adjusted Rand Index between *k*-means clustering results of cells on the DDRTree view and the known lineage labels, and **(G)** the 10-fold cross validation accuracy of a multi-class linear support vector machine (SVM) using 2D coordinates of the DDRTree view as the features and the known lineage labels as the class labels. Each comparison was repeated 100 randomly simulated data.

Fig S3

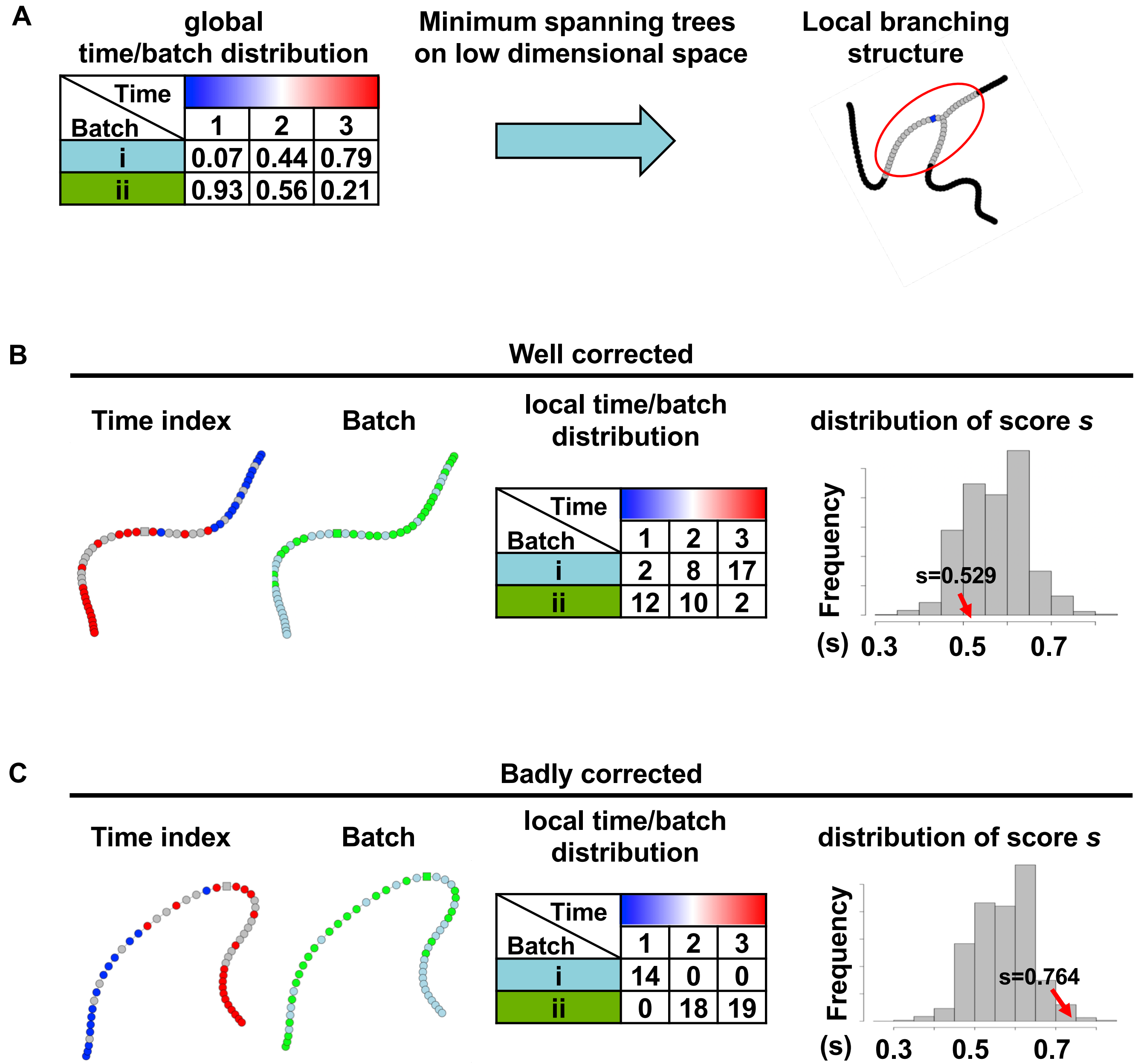
A
global
time/batch distribution
Time
Batch
1 2 3
i 0.07 0.44 0.79
ii 0.93 0.56 0.21
Minimum spanning trees
on low dimensional space
Local branching
structure
B
Well corrected
Time index
Batch
local time/batch
distribution
Time
Batch
1 2 3
i 2 8 17
ii 12 10 2
distribution of score s
Frequency
s=0.529
(s) 0.3 0.5 0.7
C
Badly corrected
Time index
Batch
local time/batch
distribution
Time
Batch
1 2 3
i 14 0 0
ii 0 18 19
distribution of score s
Frequency
s=0.764
(s) 0.3 0.5 0.7

**Fig S3. tsBET quantitatively measures the batch effects in imbalanced temporal scRNA-seq data. (A)** The real temporal scRNA-seq data usually shows an imbalance between the distribution of time indices and batch labels associated with each cell. **(B-C)** To test the batch effects in the transformed low dimensional space (e.g. the DDRTree view), the local distribution of time indices and batch labels surrounding each cell are examined at the local branching structure (e.g. the minimum spanning tree on the DDRTree view). The batch score $s$ is simply computed as the ratio of neighboring cells from the same batch. However, to account for the global imbalanced distribution between time indices and batch labels, for the same local cell populations, a null distribution of batch score $s$ is computed by using the batch labels that are randomly generated by the time indices based on the global time/batch distributions. The significance of the batch effects is then computed by comparing observed batch score $s$ and the null distribution. **(B)** In well corrected temporal data, the observed batch score $s$ is not different from the null distribution of batch score $s$, while **(C)** in the badly corrected temporal data, the observed batch score $s$ is significantly greater than the null distribution.

### Fig S4

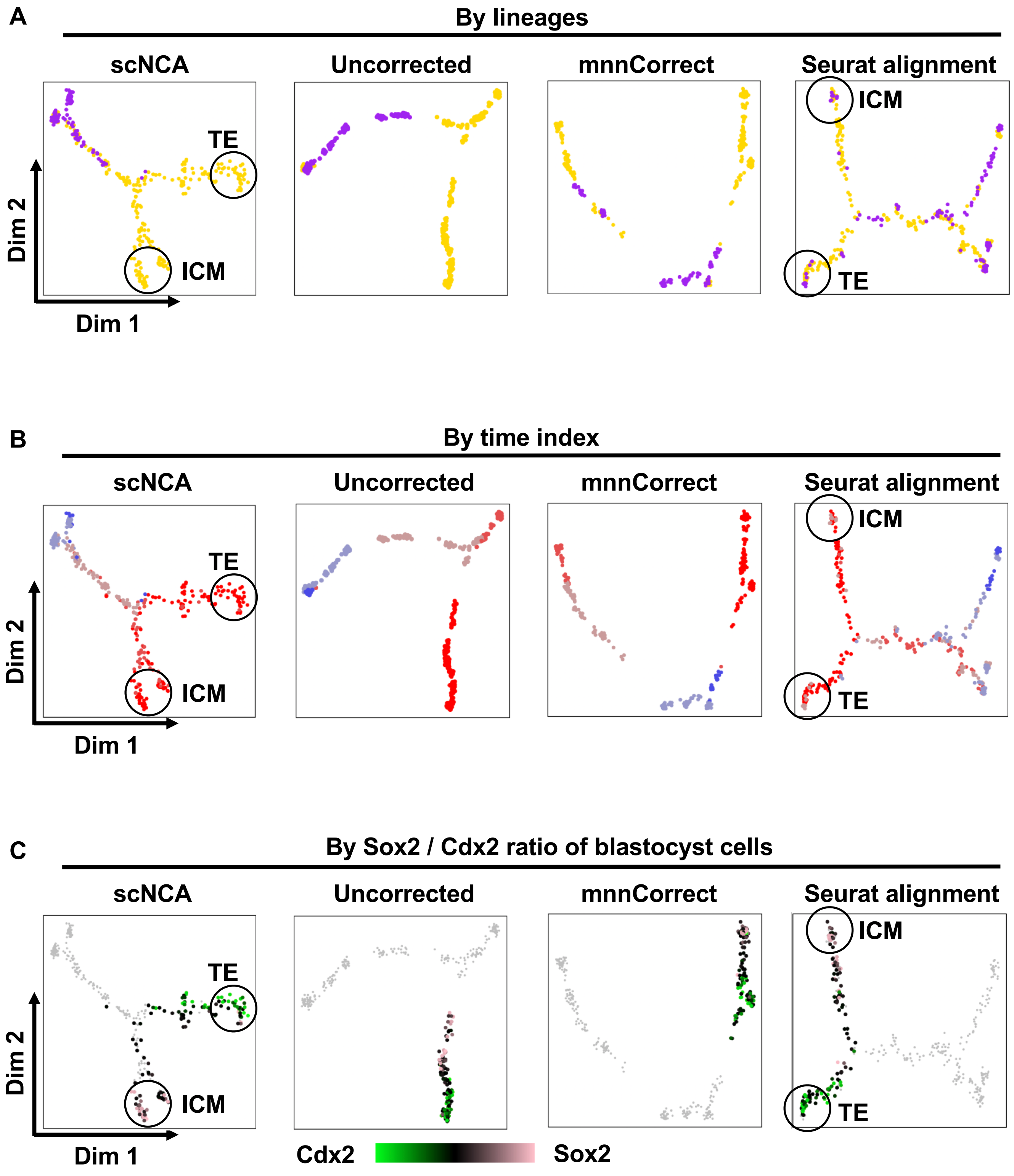


D

| Color | 2-cell | 4-cell | 8-cell | morula | blastocyst | Reference |
|---|---|---|---|---|---|---|
| (orange) | 0 | 14 | 40 | 58 | 133 | Deng |
| (purple) | 16 | 63 | 30 | 9 | 0 | Goolam |

e

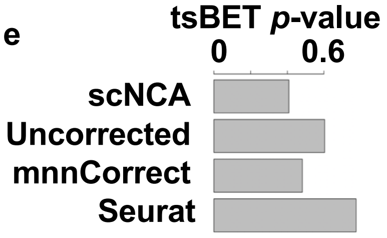

**Fig S4. scNCA integrates 354 single cells from two temporal scRNA-seq datasets during mouse preimplantation embryonic development. (A-B)** The two dimensional DDRTree view visualizes the data integrated by scNCA, mnnCorrect and Seurat alignment, as well as the uncorrected data. In panel **(A)**, the color of the points indicates the data sources, while in panel **(B)**, the color of the points indicates the associated developmental stages, from 2-cell stage (blue), to blastocyst stage (red). **(C)** The expression ratio of the TE marker Cdx2 and the ICM marker Sox2 is visualized (green: cells with relatively high Cdx2 expression levels; pink: cells with relatively high Sox2 expression levels). **(D)** The table shows the distribution of single cells across two different studies and five different developmental stages. **(E)** The tsBET for the distributions of the integrated cells on DDRTree view shows that there are no significant batch effects of the data corrected by scNCA, mnnCorrect and Seurat alignment, as well as the uncorrected data.

Fig S5

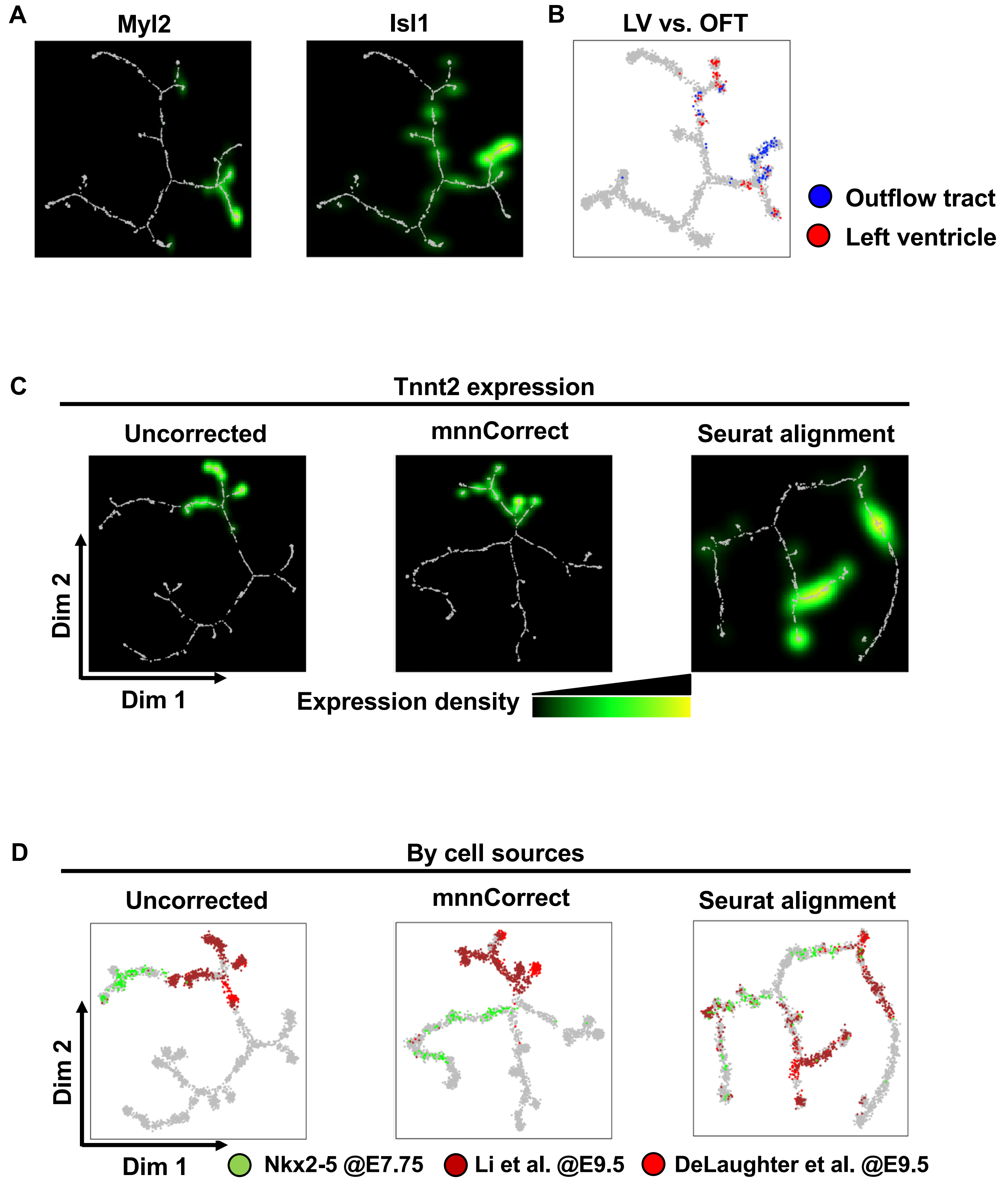
A
Myl2
Isl1
B
LV vs. OFT
Outflow tract
Left ventricle
C
Tnnt2 expression
Uncorrected
mnnCorrect
Seurat alignment
Dim 2
Dim 1
Expression density
D
By cell sources
Uncorrected
mnnCorrect
Seurat alignment
Dim 2
Dim 1
Nkx2-5 @E7.75
Li et al. @E9.5
DeLaughter et al. @E9.5

**Fig S5. The cardiovascular temporal scRNA-seq data integrated by scNCA reveal the developmental trajectory of cardiac lineages.** **(A)** The left ventricle (LV) marker Myl2 and outflow track marker Isl1 expression levels of the data integrated by scNCA indicates that the cells from cardiac lineages are enriched at the right side of the DDRTree view. The gene expression level is shown as the density of normalized read counts (black: low expression level; green: medium expression level; yellow: high expression level). **(B)** The DDRTree view indicates the whole heart cells dissected from outflow track (blue points), and whole heart cells dissected from left ventricle (red points). **(C)** The Tnnt2 expression levels are shown on the DDRTree view of the uncorrected data (left panel) and data integrated by mnnCorrect and Seurat alignment. **(D)** The DDRTree view indicates the Nkx2-5-EYFP$^{+}$ cells from E7.75 (green points), and whole heart cells from E9.5 (brown and red points), from uncorrected data, and the data integrated by mnnCorrect and Seurat alignment.

Fig S6

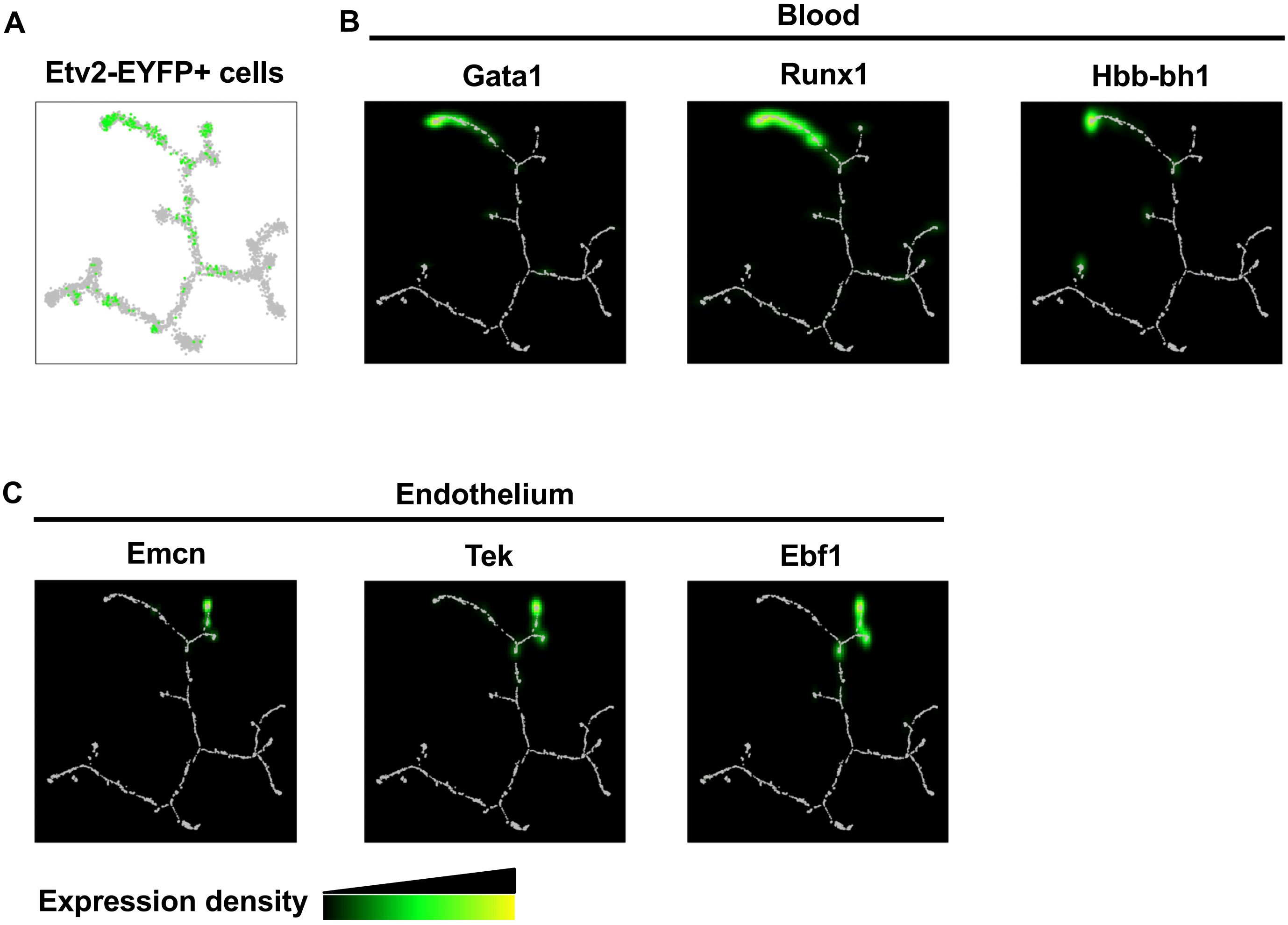
A
Etv2-EYFP+ cells
B
Blood
Gata1
Runx1
Hbb-bh1
C
Endothelium
Emcn
Tek
Ebf1
Expression density

**Fig S6. The cardiovascular temporal scRNA-seq data integrated by scNCA reveal the bifurcation of endothelial and hematopoietic lineages. (A)** The DDRTree view indicates the distribution of Etv2-EYFP$^{+}$ cells from E7.25, E7.75 and E8.25 (green points). **(B-C)** The expression levels of known **(B)** hematopoietic markers (Gata1, Runx1 and Hbb-bh1) and **(C)** endothelial markers (Emcn and Tek) shows that the cells from hematopoietic lineages are enriched at the top left branches, and the endothelial lineages are enriched at the top right branches. The gene expression level is shown as the density of normalized read counts (black: low expression level; green: medium expression level; yellow: high expression level).

### Fig S7

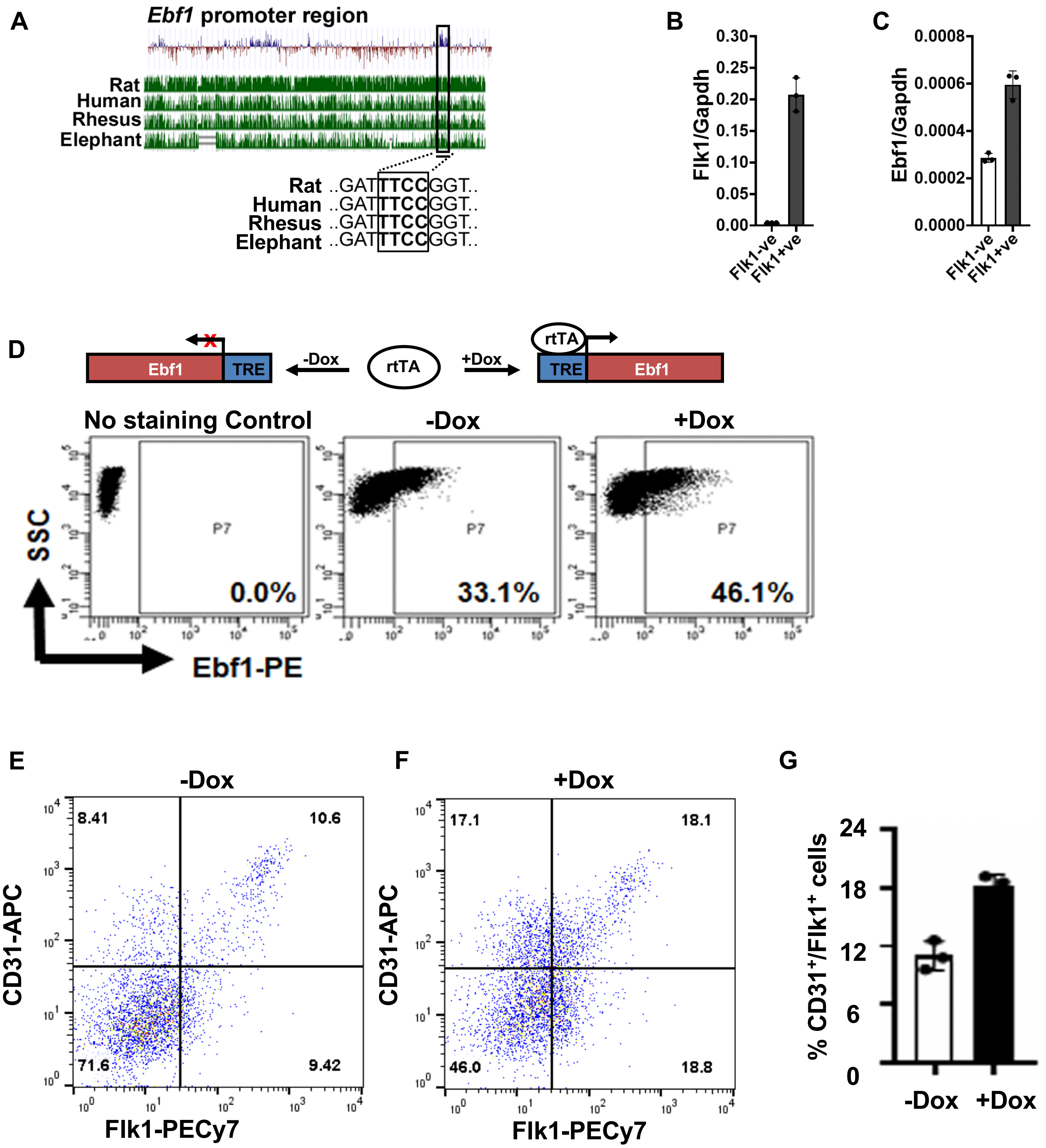

A
Ebf1 promoter region
Rat
Human
Rhesus
Elephant
Rat ..GATTTCCGGT..
Human ..GATTTCCGGT..
Rhesus ..GATTTCCGGT..
Elephant ..GATTTCCGGT..
B
Flk1/Gapdh
0.30
0.25
0.20
0.15
0.10
0.05
0.00
Flk1-ve
Flk1+ve
C
Ebf1/Gapdh
0.0008
0.0006
0.0004
0.0002
0.0000
Flk1-ve
Flk1+ve
D
Ebf1
TRE
-Dox
rtTA
+Dox
rtTA
TRE
Ebf1
No staining Control
-Dox
+Dox
SSC
P7
0.0%
33.1%
46.1%
Ebf1-PE
E
-Dox
8.41
10.6
71.6
9.42
CD31-APC
Flk1-PECy7
F
+Dox
17.1
18.1
46.0
18.8
CD31-APC
Flk1-PECy7
G
% CD31+/Flk1+ cells
24
18
12
6
0
-Dox
+Dox

**Fig S7. The Etv2 downstream target gene, *Ebf1*, promotes endothelial development.** **(A)** The UCSC genome browser track shows the evolutionary conservation of the 1.5 kb upstream fragment of the *Ebf1 gene.* The boxed region highlights the Etv2 binding motif in the promoter of the *Ebf1* gene. **(B-C)** qPCR analysis of **(B)** Flk1 and **(C)** Ebf1 from $Flk1^-$ and $Flk1^+$ sorted cells from differentiating EBs show the enrichment of Flk1 and Ebf1 transcripts in the $Flk1^+$ sorted cells. **(D)** FACS profile of Ebf1 using iEbf1 ES/EB system in -Dox and +Dox conditions. **(E-F)** FACS analysis and **(G)** quantification of endothelial lineages of endothelial markers (Flk1 and CD31) in -Dox and +Dox conditions indicate that over-expression of Ebf1 results in significant increase of the endothelial program. Error bars indicate SEM.
